# Mechanistic insights into bacterial metabolic reprogramming from omics-integrated genome-scale models

**DOI:** 10.1101/690164

**Authors:** Noushin Hadadi, Vikash Pandey, Anush Chiappino-Pepe, Marian Morales, Hector Gallart-Ayala, Florence Mehl, Julijana Ivanisevic, Vladimir Sentchilo, Jan R. van der Meer

## Abstract

Understanding the adaptive responses of individual bacterial strains is crucial for microbiome engineering approaches that introduce new functionalities into complex microbiomes, such as xenobiotic compound metabolism for soil bioremediation. Adaptation requires metabolic reprogramming of the cell, which can be captured by multi-omics, but this data remains formidably challenging to interpret and predict. Here we present a new approach that combines genome-scale metabolic modeling with transcriptomics and exometabolomics, both of which are common tools for studying dynamic population behavior. As a realistic demonstration, we developed a genome-scale model of *Pseudomonas veronii* 1YdBTEX2, a candidate bioaugmentation agent for accelerated metabolism of mono-aromatic compounds in soil microbiomes, while simultaneously collecting experimental data of *P. veronii* metabolism during growth phase transitions. Predictions of the *P. veronii* growth rates and specific metabolic processes from the integrated model closely matched experimental observations. We conclude that integrative and network-based analysis can help build predictive models that accurately capture bacterial adaptation responses. Further development and testing of such models may considerably improve the successful establishment of bacterial inoculants in more complex systems.

## INTRODUCTION

Microbiome engineering is an upcoming discipline that aims to manipulate, complement or restore the functionality of existing damaged communities, e.g., contaminated soils, by adding specific new metabolic capabilities^1^. A rational engineering approach requires a detailed understanding of general principles of the functioning of the microbial community and its physiological adaptations to perturbations, but such understanding is currently lacking and fragmentary^1,2^. The technically most simple way to provide new metabolic capacities to an existing microbial community is by strain addition (what one could call an *N+1* scenario)^3,4^. After an initial screening of the existing capacity of the microbial community, one or more preselected and well-characterized strains with the intended complementation could be prepared, formulated and inoculated into the community. Depending on the aims, such inoculants should maintain and reproduce for longer-term inside the resident community or only deploy their metabolic capacity transiently^3,4^.

Inoculation of preselected strains has been widely practised for pollutant bioaugmentation, using bacteria with particular metabolic capabilities that enable them to efficiently degrade and grow on common pollutants such as toxic aromatic compounds^5^. However, even the simplest inoculations and N+1-strategies are rarely effective because it is insufficiently understood what inoculants need to establish successfully within a (new) existing community, and how they need to adjust their physiology to meet the requirements of the new environment and degrade the desired toxic compound(s). Modeling strategies based on the integration of a variety of (nowadays more easily) accessible condition-specific omics data, would help to better understand and predict how cellular regulation and physiology at different growth conditions and environments interplay. However, the impact and advantage of such integrative analysis are not yet explored to its full extent^6^. We propose and demonstrate here that combining comprehensive genome-wide transcriptomics, exometabolomics and metabolic modeling can better predict physiological adaptation.

Metabolic modeling has largely advanced through the development of GEnome-scale Metabolic models (GEMs) and constraint-based modeling techniques such as Flux Balance Analysis (FBA). GEMs can be built from the annotated genomes and they describe an organism’s metabolism as completely as possible, linking genotype to metabolic phenotypes^7^. GEMs encompass metabolites, metabolic reactions, and genes coding for the enzymes catalyzing the reactions. Together with FBA, GEMs predict steady-state fluxes^8,9^, and therefore, they can predict cellular physiology. While the genome specifies the complete set of biochemical reactions which the cell can potentially carry out, the actual enzymatic capacity at each physiological condition is orchestrated by regulatory networks in the cell. GEMs do not explicitly consider regulation, whose effects are better reflected in the global transcriptome and the metabolome^10–14^. FBA approaches have been extended with RNAseq and metabolomics data to capture cell regulation and more accurately describe cellular metabolic behavior^15^. For example, transcriptional regulation of gene expression has been linked to GEMs, either by taking into account the absolute expression values, scoring genes and subsequently reaction fluxes as active or non-active based on their expression,^16–18^ or by incorporating relative gene-expression^14,18^. Use of relative gene expression is assuming that the relative changes between two conditions correlate with the resulting differential flux profiles. Both approaches can lead to condition-specific GEMs that are more effective for inferring the actual biochemical activity and the observed physiology of the microorganism.

As a study system for predicting physiology from an integrated GEM-transcriptome-metabolome approach, we here use *Pseudomonas veronii* 1YdBTEX2. Strain 1YdBTEX2 is capable of degrading a variety of mono-aromatic hydrocarbons such as benzene, toluene, ethylbenzene, and *m-* and *p-*xylene (BTEX)^21–23^. The ability of *P. veronii* 1YdBTEX2 to grow in contaminated environments makes it a promising candidate for rational complementation of microbial communities in contaminated soils^24^. Based on an available manually curated high-quality genome^23^, we reconstructed the first GEM for *P. veronii* (iPsvr). Genome-wide transcription changes and exometabolome compounds were measured during growth of *P. veronii* on toluene, in exponential and in stationary phase. Transcriptome and exometabolome data were integrated into the iPsvr using the recently developed tool *REMI* (<u>R</u>elative <u>E</u>xpression and <u>M</u>etabolomics Integrations)^20^. Two obtained metabolic models representing exponential and stationary physiologies were then used to evaluate growth rates and the production of biomass precursors, and model predictions were compared to the experimentally observed values. Although the temporal variations of the growth rate cannot be predicted using GEMs^25^, we showed that introducing the additional regulatory information from gene expression and metabolomics data into GEMs allows for consistently estimating growth rates at different growth phases. Finally, we incorporated into iPsvr previously published transcriptomics data of *P. veronii* transits from liquid culture to sand^23^ to understand its physiological adaptation in soil. Our work shows strong consistency of model outputs with the experimental data, manifesting that integration of condition-specific omics data into a curated GEM constitutes a major improvement for prediction of metabolic reprogramming during adaptation.

## RESULTS

### Developing an integrated genomic-transcriptomic-metabolomic workflow

To develop a pipeline that integrates genomics with transcriptomic and metabolomic data we advanced in three stages: 1.) Quantify the cellular states at each unique growth phase by genome-wide transcriptomics, and exometabolomic data from spent media composition (Fig. 1A); 2.) Construct a GEM for *P. veronii* strain 1YdBTEX2 (iPsvr), gap-fill missing parts of the metabolism (compounds and reactions), complement genome annotation using the transcriptomics and exometabolomic data and estimate the steady-state growth rate using FBA (Fig. 1B); and 3.) Link the interrelationships between growth phases and the differentially expressed genes and metabolite abundances by statistical inference and by REMI. The pipeline generated two growth-phase-specific models, iPsvr-EXPO and iPsvr-STAT (Fig. 1C), which were used to predict quantitative and dynamic readouts of *P. veronii* metabolism in both conditions and in liquid-to-sand transition.

**Figure 1:**
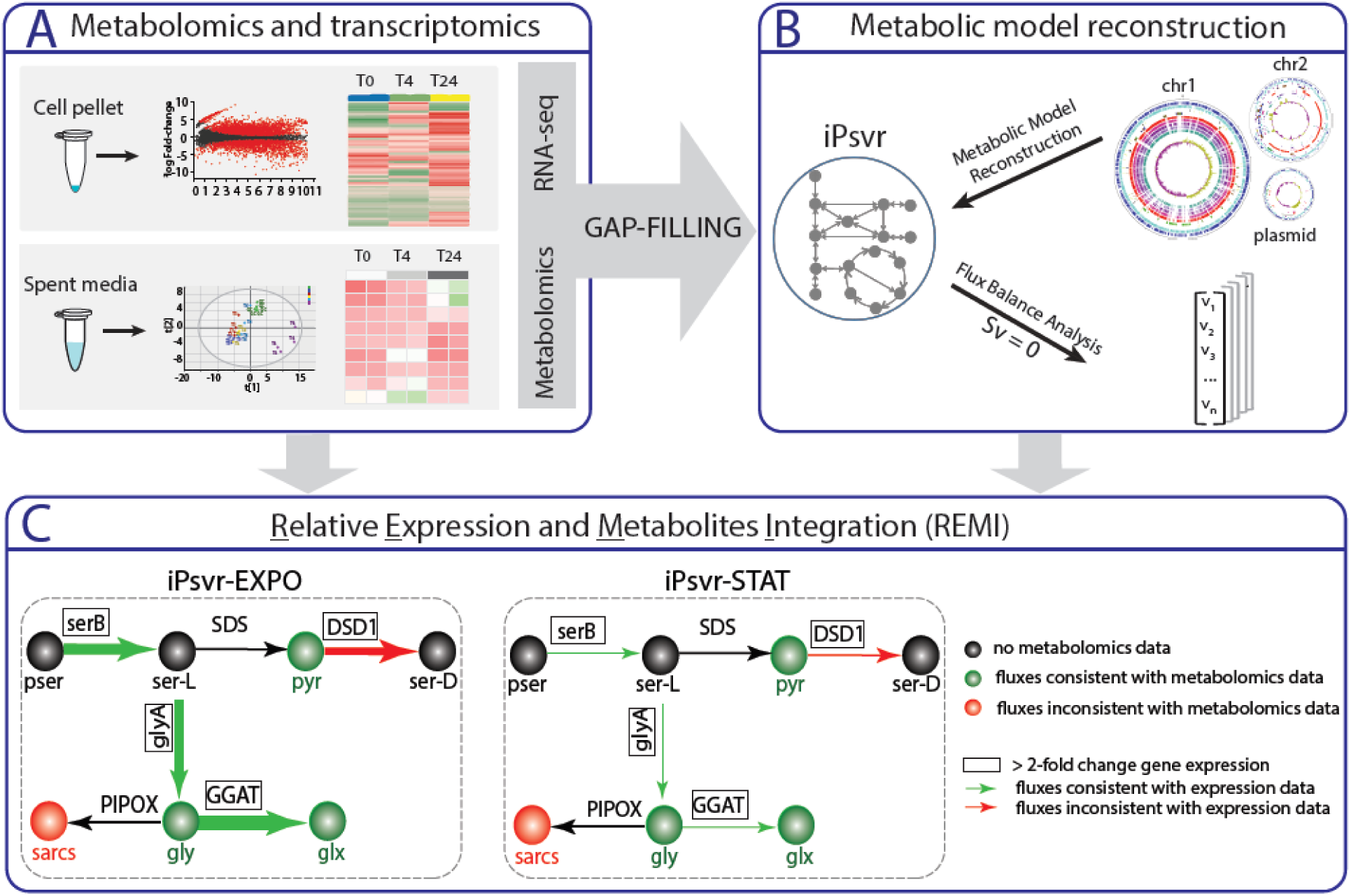
Schematic overview of the integrated genomic-transcriptomic-metabolomic pipeline applied in this study. (A) Stage 1: Relative gene expression and exometabolomic data were determined and analyzed, and these data were used to gap-fill iPsvr at blocked reactions. (B) Stage 2: A genome-scale metabolic model (GEM) of *P. veronii* strain 1YdBTEX2, iPsvr, was reconstructed and flux-balance analysis (FBA) was performed to simulate the growth of the cell. (C) Stage 3: Additional relative differential gene expression and metabolite abundance data were integrated into the metabolic model with REMI and physiology-specific models were built. Here, the REMI methodology is illustrated on a section of inferred iPsvr glycine, serine and threonine metabolism and iPsvr-EXPO (exponential phase) and iPsvr-STAT (stationary phase) as the two physiology-specific models. Significantly differentially expressed genes (here, *serB*, *glyA*, DSD1 and GGAT) are outlined in boxes. The thickness of arrows designates the fold-change in estimated fluxes, where green arrows indicate consistency with the gene-expression fold-change values, and the red ones inconsistencies. Measured metabolite concentrations (here: pyr, sarcs, gly and glx) are indicated in green if the values are consistent with estimated fluxes and in red otherwise. Phosphoserine phosphatase, SerB; Serine hydroxymethyltransferase, GlyA; D-serine dehydratase, DSD1; Glyoxylate aminotransferase, GGAT; Pyruvate, pyr; Sarcosine, sarcs; Glycine, gly; Glyoxylate, glx.

### Genome-wide gene expression and metabolite formation over time

Whole-genome gene expression profiles and metabolite formation in the spent medium were analyzed in *P. veronii* cultures growing in liquid minimal medium with toluene as sole carbon and energy source, sampled at 0 h (T0h), 4 h (T4h, EXPO) and 24 h (T24h, STAT) after inoculation. Genome-wide gene expression was quantified by mapping Illumina 100 nucleotide long single–end sequencing reads from deeply sequenced cDNA libraries to the protein coding genes in *P. veronii* genome (read numbers indicated in Table S1). For each sampling time point, four replicates clustered closely together, with slightly higher variability observed among the T24h replicates (Fig. S1A). A pair-wise comparison of expression levels showed that 1458 (818 up-regulated and 640 down-regulated) out of the total 6943 genes (21%) were significantly differentially expressed between EXPO and STAT phase cells, with at least 2-log fold-change induction (false discovery rate [FDR]<0.05) (Fig. 2A, B).

**Figure 2:**
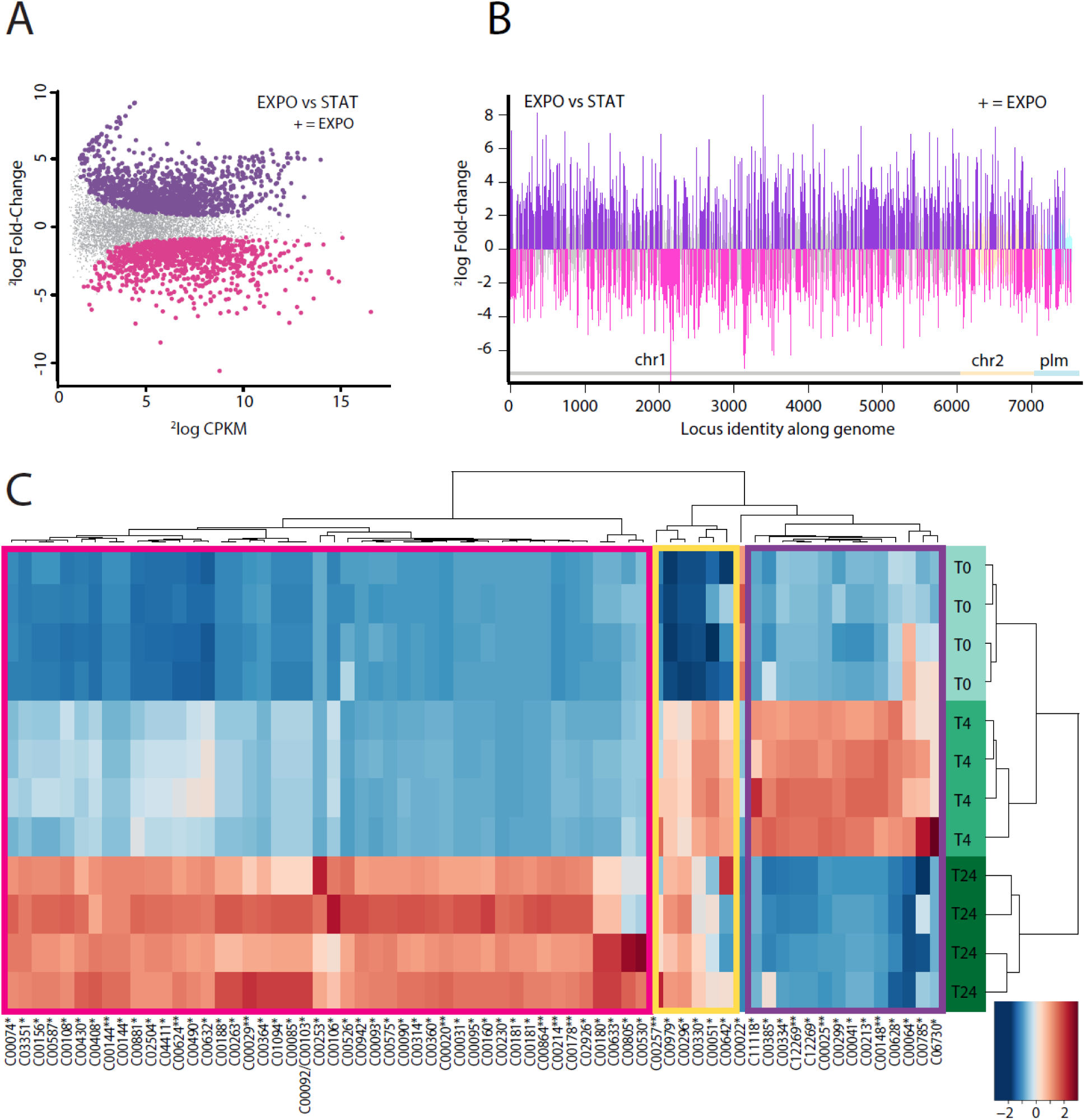
Genome-wide gene-expression and metabolite abundance differences in *P. veronii* 1YdBTEX in exponential (EXPO) relative to stationary (STAT) phase. (A) Smear-plot of global gene expression intensity (2log counts per kilobase per million, CPKM) versus expression changes (2log fold change) in EXPO vs. STAT. In grey, genes not statistically differentially expressed (logFC<1, FDR>0.05, P>0.01); magenta, genes with lower, and dark purple, genes with higher expression in EXPO (+). (B) Differential expression per gene between EXPO and STAT plotted as a function of genomic location (chromosome 1, chr1; chromosome 2, chr2; and plasmid, plm; organized according to locus_tag number). Bars indicate 2log-fold change, dark purple and pink denote statistically significant higher and lower expressed genes in EXPO, respectively. (C) Heatmap of 164 metabolites annotated with their KEGG IDs showing clustering at different culture sampling time points. Metabolites are clustered (x-axis) in three major groups (left to right): (i) accumulation at T24h (STAT) (pink box), (ii) accumulation at T4 continued to T24 (yellow box), and (iii) accumulation at T4h (EXPO) followed at depletion at T24h (STAT) (purple box).

*A priori*, based on the genome annotation, a subset of 1241 “metabolic genes” were used in the GEM reconstruction (iPsvr). Out of these 1241, 300 (21%) were significantly differentially expressed in EXPO vs. STAT phase cells (FDR<0.05) (Table S2). The transition to STAT phase in bacteria is characterized by growth arrest in response to several factors, such as nutrient depletion, the accumulation of toxic compounds and environmental stress, which decrease ribosomal activity and therefore protein synthesis. As anticipated, enriched GO terms for the category “Biological Process” among the differentially expressed genes between EXPO and STAT included “protein folding” (GO:0006457), “tRNA aminoacylation for protein translation” (GO:0006418), “intracellular protein transmembrane transport” (GO:0065002) and “regulation of transcription, DNA-templated” (GO:0006355) (Table S3), thus indicating cells to be more active in EXPO phase, as expected. Consistent with nutrients becoming depleted in STAT phase, the terms “benzoate catabolic process via hydroxylation” (GO:0043640) and “tricarboxylic acid cycle” (GO:0006099) (Table S3), important for aromatic compound catabolism, were under-represented in the STAT phase transcriptome.

The untargeted metabolomic analyses of the spent medium detected 1630 (positively charged) and 3509 (negatively charged) distinct ion species or metabolite features. Unsupervised principal component analysis yielded three distinct clusters indicating metabolic phenotype differentiation over time, from inoculum to stationary phase (Fig. S1B). Similar to the transcriptomics data, a greater variability was observed among the T24h replicates.

Temporal patterns of annotated metabolites by HMDB database (accurate mass) matching^26^ showed a significant increase in the spent media over time of the majority of the metabolites implicated in the toluene and benzene degradation pathways and central carbon pathways, including glycolysis, purine and pyrimidine metabolism and amino acid metabolism (Fig. 2C). This implies their production by the bacteria and progressive release into the media. One specific group of metabolites, including citrate (C00158), glutamate (C00025), glutamine (C00064), and aspartate (C00049), accumulated significantly in EXPO phase (T4h) in comparison to T0h, followed by lower levels in STAT phase (Fig. 2C). This suggests their excretion in EXPO phase and subsequent reconsumption when other nutrients became limiting (Fig. 2C).

### Genome-scale metabolic model (GEM) of *P. veronii* strain 1YdBTEX2 (iPsvr)

A draft GEM was generated from the curated *P. veronii* genome^23^ by using the RAVEN toolbox^27^ (Fig. 1B). The draft GEM was gap-filled by iterative manual curation until we obtained a model able to carry non-zero flux through the biomass reaction at steady state. This signified cell ‘growth’, and indicated that the model was performing as expected for a biological system. The cell biomass composition and compartment information were derived from two available models of other Pseudomonads species: *P. putida*^28–30^ and *P. stutzeri*^31^ (see Methods). The reconstructed iPsvr accounted for 1243 genes, 1812 metabolic reactions and 1677 metabolites localized within two intracellular compartments, the cytosol and periplasm, and the extracellular environment (Table 1).

**Table 1:**
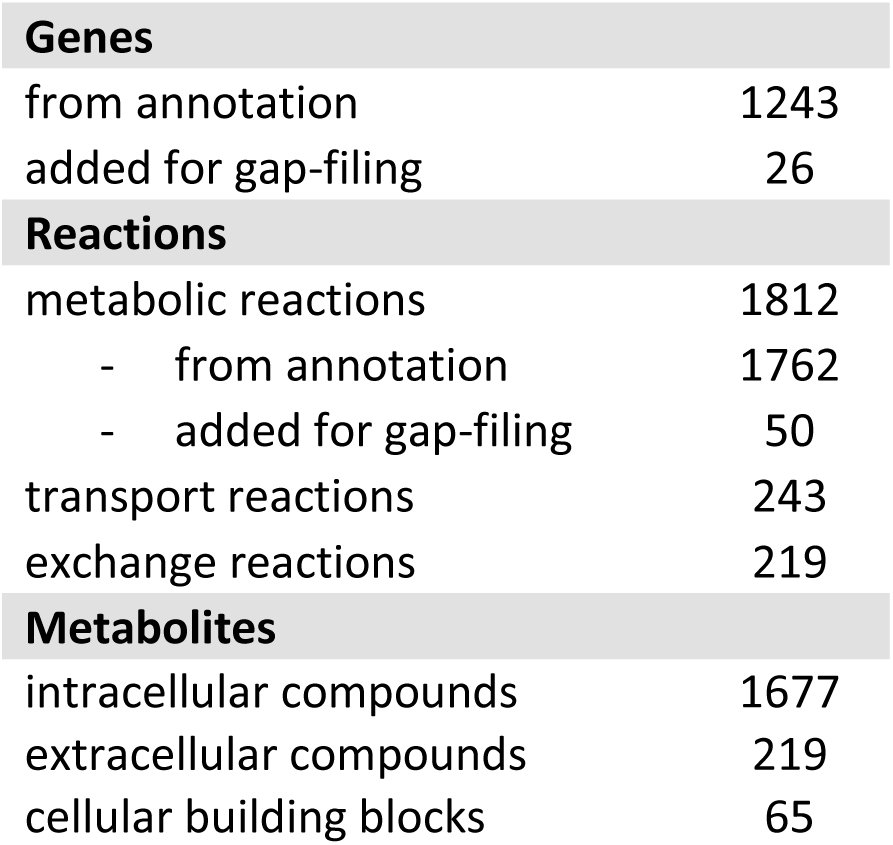
iPsvr components in the final GEM.

The scope of iPsvr GEM was further widened by restoring the connectivity of the remaining ‘blocked’ reactions, i.e., isolated reactions that carry zero flux at any condition. To this end, we explicitly considered the empirical gene-expression and exometabolomics data. We first used a graph-based algorithm (see Methods) to decompose the iPsvr metabolic network into its main subnetworks of 1370 reactions and 23 blocked reactions/pathways of different lengths, with the longest blocked pathway consisting of seven reaction steps (Table S4). Out of 191 blocked reactions/pathways, we identified those associated with differentially expressed genes between the two growth conditions and the ones whose participating metabolites were present in the exometabolomic data. The identified reactions/pathways were next unblocked by gap-filling as described in the Methods section. Interestingly, we identified gap-filling reactions that had been annotated to *P. veronii* genes with RAVEN but had a lower score than the ones chosen as a baseline for the draft reconstruction of iPsvr. The gap-filling algorithm introduced 50 new metabolic reactions together with their corresponding 26 genes to iPsvr (Table 1 and Table S5).

The experimentally determined maximum specific growth rate of *P. veronii* strain 1YdBTEX2 in minimal medium with toluene as the sole carbon source was in the range of 0.25 h^-1^ to 0.35 h^-1^. Computational prediction of the growth rate on toluene from the curated iPsvr using thermodynamics-based flux balance analysis (TFA)^8,32,33^, which integrates thermodynamic constraints into FBA, yielded 0.91 h^−1^ (at a maximum allowed toluene uptake rate of of 5.5 mmol gDW^−1^ h^−1^, and without considering transcriptomics and metabolomics information, see Methods section).

### Omics-based curation and gap-filling in iPsvr for toluene degradation and phenylalanine metabolism

Given the importance of toluene degradation by *P. veronii* 1YdBTEX2, we manually curated the predicted toluene (Fig. 3A) and phenylalanine metabolic pathways (Fig. 3B) to ensure they were fully functional in iPsvr. Toluene is converted in *P. veronii* via (1S,2R)-3-methylcyclohexa-3,5-diene-1,2-diol to 3-methylcatechol, which is further degraded, according to the KEGG pathway database^34^, through two pathways until the central carbon metabolism compounds are reached (Fig. 3A and Fig. S2). The iPsvr growth simulation on minimal media containing toluene as the sole carbon source showed that the pathway producing pyruvate and acetaldehyde in four reaction steps was functional (1.13.11.2, 3.7.1.-, 4.2.1.8 and 4.1.3.39 in Fig. S2), which has experimentally been shown to be the main toluene degradation route in *P. veronii*^23^. In contrast, the second functional pathway (Fig. 3A)^23^, was blocked in iPsvr because it lacked the enzyme 2.8.3.6 (highlighted in red in Fig. 3A), and it was therefore disconnected from the main metabolic subnetwork. We found that three out of the six genes in this pathway (PVR_r1g5041, PVR_r1g5042, PVR_r1g1440) were more than 2-fold differentially expressed between the two growth conditions (highlighted in green in Fig. 3A), which further suggested that this pathway is indeed active in *P. veronii*. Homology-based BLAST searches^35^ of the gene sequences of 2.8.3.6 against the *P. veronii* genome identified the corresponding gene for catalyzing this reaction (PVE_r1g3867, e-value of 10^-20^, to *scoA*). Note that the default e-value in RAVEN is 10^-50^, which is why the reaction was not initially captured in the model from the genome annotation. Therefore, the missing reaction (2.8.3.6) was added to iPsvr, and the toluene pathway (Fig. 3A) was connected to the rest of the metabolic network through the Krebs cycle and carried flux.

**Figure 3:**
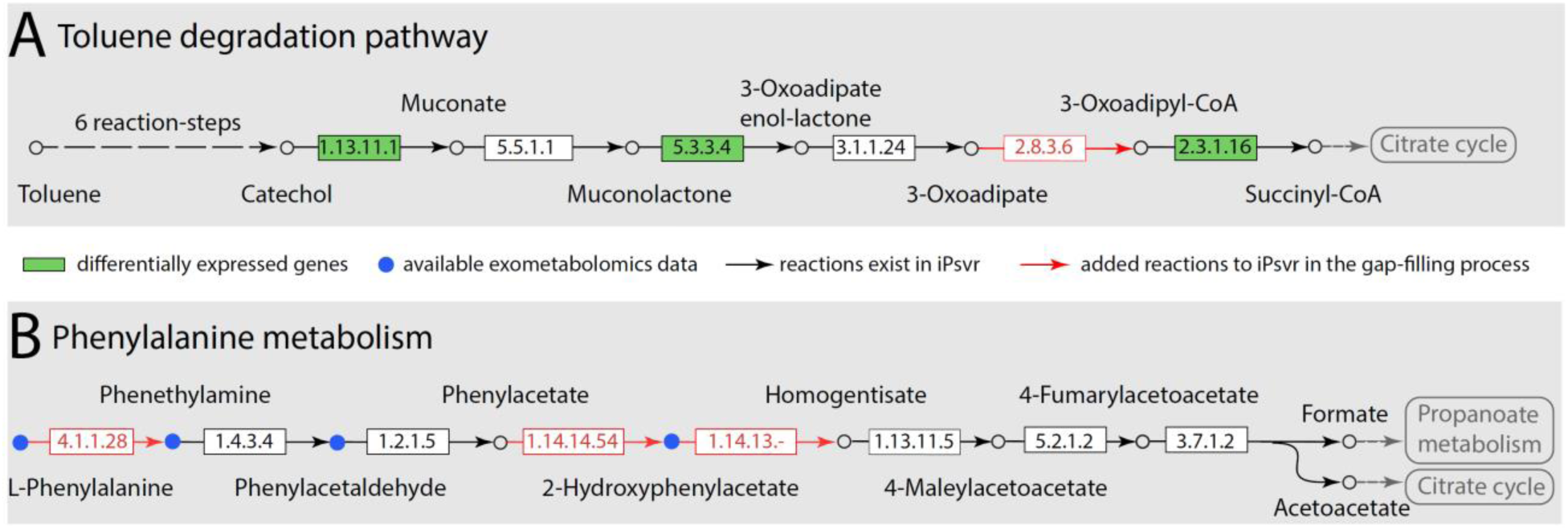
Gap filling of two blocked pathways in iPsvr based on the gene expression and exometabolomics data. (A) One of the two toluene degradation pathways in *P. veronii* (from the KEGG pathway database^34^) involved one missing enzyme (highlighted in red), and three genes (highlighted in green) differentially expressed between the exponential and stationary growth conditions. (B) The phenylalanine metabolic pathway involved three missing enzymes (highlighted in red) and four metabolites (highlighted in blue) identified in the exometabolomic data.

The phenylalanine metabolic pathway was gap-filled using the exometabolomic data (Fig. 3B). Three out of the eight reaction steps of this pathway (4.1.1.28, 1.14.14.54 and 1.14.13.-) were initially missing in iPsvr, leading to a dead-end pathway without flux. Four metabolites of phenylalanine metabolism were detected in the exometabolomics data (colored in blue in Fig. 2B), one of which, 2-hydroxyphenylacetate, was absent in the iPsvr GEM. This suggested that phenylalanine metabolism should proceed via 2-hydroxyphenylacetate in *P. veronii* (Fig. 3B). The enzyme producing 2-hydroxyphenylacetate from phenylacetaldehyde (1.14.14.54) was found by BLAST (PVE_r1g94, e-value of 10^-20^ to CYP504), added to iPsvr and could thus link both metabolites. The next step (1.14.13.-), yielding homogentisate (Fig. 3B) was also added, but no protein sequence could be assigned, as it corresponds to an orphan reaction (KEGG reaction ID: R05450). The first enzyme (4.1.1.28), which decarboxylates L-phenylalanine to phenethylamine, must be present, given the detection of both compounds in the exometabolomics data, although no *P. veronii* protein sequence could be identified through BLAST search.

### Omics integration into iPsvr

REMI (<u>R</u>elative <u>E</u>xpression and <u>M</u>etabolomics Integrations) was used to integrate the relative gene-expression and metabolite abundance data between EXPO and STAT into iPsvr. We explored three scenarios, depending on the type of data integrated into iPsvr: (i) REMI-TGex for the integration of relative gene-expression data only; (ii) REMI-TM for the integration of metabolomics data only; and (iii) REMI-TGexM for the simultaneous integration of both the gene-expression and metabolite abundance data into a thermodynamically curated model. The “T” in the three methods stands for the inclusion of thermodynamic constraints. Contextualized metabolic models (iPsvr-EXPO and iPsvr-STAT) coherent with omics data were built, and the growth rates were again simulated (Table 2).

**Table 2:** Summary of the REMI results of the relative integration of exometabolomics (REMI-TM), gene expression (REMI-TGeX) and both datasets (REMI-TGeXM) into iPsvr. Theoretical maximum consistency score, TMCS; Maximum consistency score, MCS; Std, standard deviation.

| Method | Score | | Growth rate ($\text{h}^{-1}$) | |
| --- | --- | --- | --- | --- |
|  | TMCS | MCS | EXPO | STAT |
| TFA | - | - | $0.91^*$ | $0.91^*$ |
| REMI-TM | 215 | 123 | 0.86 | 0.67 |
| REMI-TGex | 618 | 229 | 0.48 | 0.06 |
| REMI-TGexM | 833 | 342 | 0.40 | 0.07 |
| Experimental | - | - | $0.31 \text{ (std=0.05)}$ | 0 |

*\*by definition, there is no difference between the simulated growth rate at different growth phases in FBA simulations.*

342 out of a total of 352 (97%) integratable gene-expression and metabolite abundance data points could be consistently included in REMI; the remaining 3% being inconsistent (Table 2, third row). Simulated growth rates for EXPO phase under the condition of integrating both gene-expression and metabolomics data (REMI-TGexM) were the closest to the experimentally observed rates (0.40 h^-1^ for REMI-TGexM versus 0.25–0.35 h^-1^, Table 2).

The integration of exometabolomics data alone (REMI-TM) was insufficient, with a predicted growth rate of 0.86 h^-1^, which is close to the rate of 0.91 h^-1^ that was obtained for the unconstrained iPsvr model (Table 2). The integration of relative gene-expression data alone (TGex) yielded an *in silico* exponential growth rate of 0.48 h^-1^, which is close to the REMI-TGexM value (Table 2). Both REMI-TGex and REMI-TGexM, but not REMI-TM, correctly estimated near zero growth rates (0.06 h^-1^ and 0.07 h^-1^, respectively) for stationary phase cells (Table 2).

### Analysis of the biomass precursor production at different growth phases

To understand the underlying mechanisms of growth reduction when cells transit from EXPO to STAT, we identified the biomass precursors that may become limiting in the stationary phase, therefore leading to growth arrest. The 65 biomass precursors were grouped into seven groups of biomass building blocks (BBBs), namely carbohydrates, cofactors and vitamins, DNA nucleotides, lipids, minerals, amino acids, and RNA nucleotides. For each metabolite, we calculated the log2 fold-change of their maximal production between EXPO and STAT using REMI-TGex and REMI-TGexM (Table S6).

Both REMI-TGex and REMI-TGexM indicated large variations in the production of BBBs between the two phases (Fig. 4), the maximal relative change occurring in the production of cofactors and vitamins. Out of the 20 precursors that were classified as cofactors and vitamins, the production of 10 (REMI-TGexM) and 18 (REMI-TGex) of them was at least 2-log fold-change higher in iPsvr-EXPO than in iPsvr-STAT (Table S6). Rather surprisingly, an increased production of certain BBBs occurred in STAT. Both REMI-TGex and REMI-TgexM predicted that several amino acids, e.g., tyrosine and lysine, and some lipids, e.g., hexadecanoic acid, had at least a 2-log fold-change higher production in STAT (Fig. 4A & B). As before, the variant with integration of only exometabolomic data (REMI-TM) did not predict any significant differences in the production of BBBs between EXPO and STAT culture (Table S6).

**Figure 4:**
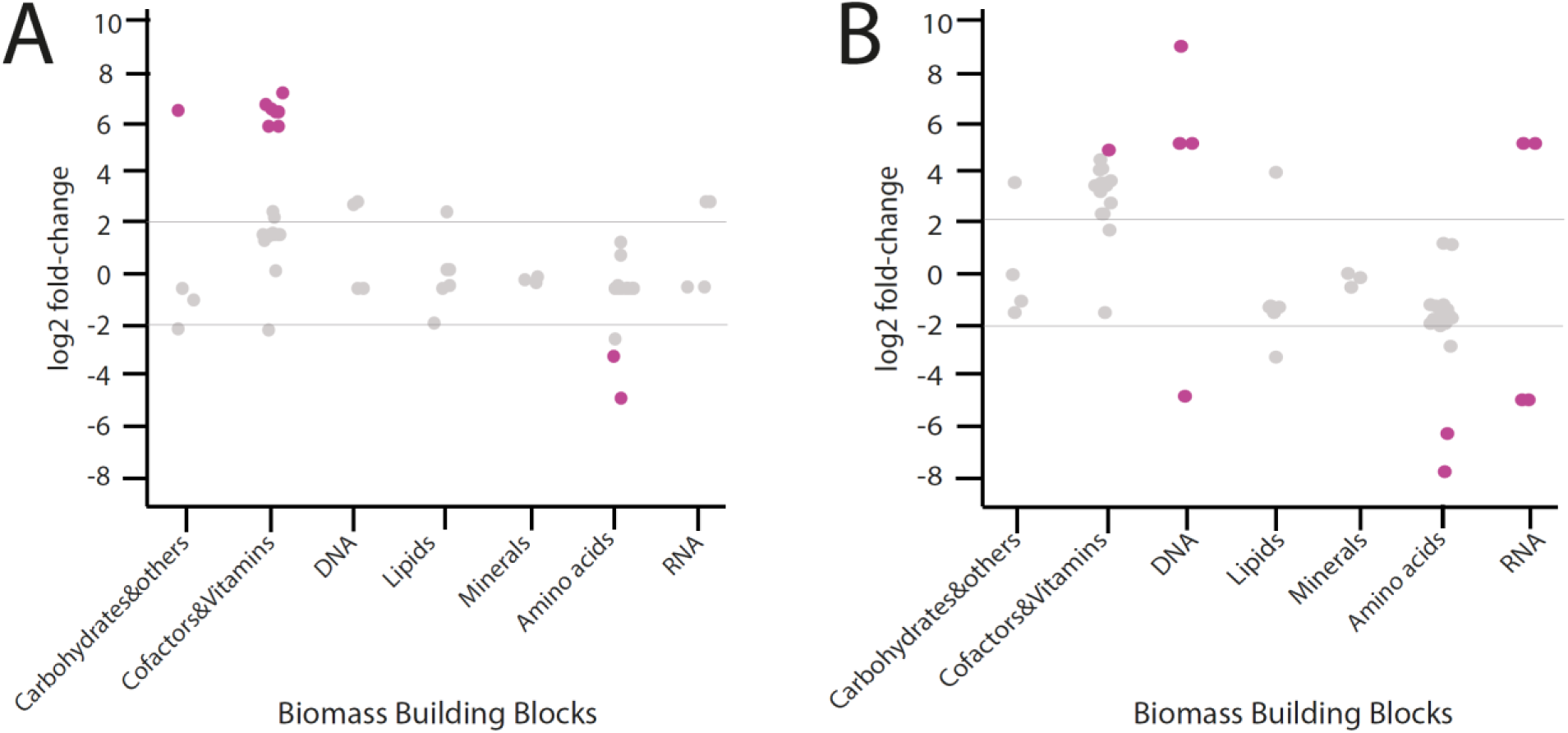
Differential *in silico* production (mmol/gDW/h) of seven biomass building block groups between the exponential and stationary growth phase. (A) Integration of relative gene expression data (EXPO vs STAT) using REMI-TGexM. (B) Integration of relative gene expression and exometabolomics data (EXPO vs STAT) using REMI_TGex. Each dot in the graph represents the individual biomass precursors within that category and the statistically significant changes (p=value < 0.05) are highlighted in purple. For BBB group statistics, see Table S7.

### *P. veronii* adaptation from liquid culture to the soil as growth environment

Finally, as an additional validation of our approach, we predicted the *in silico* physiology for *P. veronii* during adaptation to the soil environment, using a previously published genome-wide transcriptome data set of cells exposed for 1 h to liquid medium or to sand, with either toluene or succinate as carbon substrate^23^.

Integrating the transcriptomic data^23^ into iPsvr using REMI-TGex, produced *in silico* growth rates of 0.52 and 0.66 h^−1^ for toluene or succinate in liquid medium. Remarkably, the model predicted strong reduction of growth rate upon transition to the sand, i.e., from 0.52 h^-1^ to 0.23 h^-1^ (toluene) and 0.66 h^-1^ to 0.07 h^-1^ (succinate). This suggests that cells have to adapt to soil as their new environment and have to reprogram their physiology before resuming growth. The higher reduction of growth in case of succinate is in agreement with experimental results in the previous study^23^, and can be explained from the lower concentration of succinate than toluene available in the soil, therefore rapidly used by the cells and leading to nutrient starvation.

To understand how the transition from liquid medium to sand constrains the metabolic fluxes and impacts growth of *P. veronii*, we analyzed the production of BBBs in both growth environments. Intrestingly, we observed the same robust reduction in the production of cofactors and vitamins upon transition to sand in the case of succinate, as had been predicted for transition to STAT in liquid culture (14 out of 20 precursors with more than 2-log fold lower production). In contrast, in case of cells exposed to toluene in sand only 8 of the cofactor and vitamin precursors were 2-log fold lower expressed. The higher number of precursors with lower production rates might thus explain the stronger growth arrest in case of succinate during transition to sand as opposed to toluene. In contrast, the production of other precursor BBBs, such as phosphatidylcholine and ubiquinone-8, similarly increased in sand both for toluene and succinate (Fig. S3 A&B). “This suggests that the cells in sand reshape their membrane content and respiratory pathways

## DISCUSSION

This work demonstrates how a metabolic-model-based (multi)omics data integration approach can accurately capture cell physiology and adaptation during growth or environmental transitions. Metabolic models are typically used for predicting optimal cell growth, but not for growth transitions and adaptation. We show here the usefulness of models with comprehensively integrated transcriptomics and exometabolomic to capture changes in growth rate during adaptation. As example, we studied transition of the toluene-degrading bacterium *P. veronii* 1YdBTEX2 from exponential growth to stationary phase, and during adaptation from liquid to a sand environment.

For the purpose of this study, we reconstructed and curated the first genome-scale metabolic model of *P. veronii*, iPsvr. iPsvr is useful for a wide range of applications in biological and biotechnological research, such as the integration of omics data, investigation of relationships among species in microbial communities, bioremediation strategies, design of metabolic engineering strategies, hypothesis-driven discovery, and analysis of metabolic network properties. As such, iPsvr may represent a valuable resource for the study of complex microbiomes and microbiome engineering.

GEMs usually contain inconsistencies that manifest as blocked reactions, i.e., the reactions that carry zero flux. Most of the available metabolic models are gap-filled to obtain a functional model that produces all the BBBs. Nevertheless, many reactions in GEMs are not directly connected to biomass formation and may remain blocked even after routine gap-filling. Roughly, 20–50% of the reactions of published GEMs are blocked reactions^36,37^. For example, in the most recently and best-curated published *E. coli* model, iML1515, 10% of the reactions are blocked^38^. We identified the blocked reactions/pathways in iPsvr and introduced a novel omics-based gap-filling approach, which targets the gap-filling of reactions/pathways by looking at differentially expressed genes or metabolites. We showed that the gap-filling step improved the metabolic model connectivity, thus increasing the number of functional pathways that can carry flux. The curation also increased the consistency of the metabolic model with the omics data, expressed as the number of genes/metabolites with available omics data that can be integrated into iPsvr. We illustrated through two gap-filling examples on toluene and phenylalanine metabolism how we could add the missing parts of metabolism and increase consistency.

Statistical analysis of gene expressions and metabolite levels revealed significant variations across the exponential and stationary phase, exemplifying the known distinct physiologies under these conditions. However, the system’s level physiology and the underlying adaptation mechanisms remained unrevealed.

It should be highlighted that in FBA, the growth rate is evaluated under the assumption that all metabolic fluxes in the cell are geared towards the maximal production of biomass at each moment of the cell growth and under any environmental condition. Nevertheless, inclusion of REMI^20^, a method for the integration of omics data into metabolic models and for building context-specific models, produced more accurate predictions of *P. veronii* growth rates under dynamic conditions. Integration of the transcriptomic data (as in REMI-TGex and –TGexM) proved to be vital for improving growth rate predictions, whereas inclusion of metabolomic data alone (as in REMI-TM) was insufficient. This might be expected, because only a small subset of metabolites could be putatively identified, such that their impact on the predictions was comparatively smaller. The estimated growth rate was slightly closer to the experimentally measured values in the case of REMI-TGexM, which combines transcriptomics and metabolomics data. Overall, these results clearly demonstrated that integrating multi-omics data in GEMs significantly increases the consistency of the model predictions with experimental observations.

REMI-iPsvr analysis suggested *P. veronii* adapts to STAT phases primarily by limiting the metabolic fluxes for the production of the majority of cofactors and vitamins required for growth, with metabolism reshuffling being regulated at the transcriptional level. In addition, an increased production of specific amino acids in STAT phase suggests the cell is preparing for starvation and survival. Similar to the EXPO to STAT transition, the production of most of the BBBs decreased in cells inoculated in sand compared to liquid culture, which explains the overall observed reduction in growth rates. In contrast, the production of several precursors from the *cofactors and vitamins*, and the BBBs *amino acids* actually increased in cells transited to sand, both for sand with toluene or succinate. We hypothesize that these BBBs are needed to adapt to the local sand conditions. This finding is in agreement with work of Morales et al.^23^, who concluded from Gene Ontology terminology that *P. veronii* cells inoculated in sand readjusted their metabolism during the first hour of contact.

Collectively, our results demonstrate the importance of integrating into metabolic models the contextualization of condition-specific gene-expression and metabolite-abundance data. This increases the value of growth rate predictions and improves the assessment of the relative changes of measurable metabolic phenotypes. The method as demonstrated here is thus an important advancement to explain, quantify or predict cellular responses to environmental or genetic perturbations, which is crucial for microbiome engineering.

## METHODS

### Culture conditions for *P. veronii* transcriptome and exometabolomic studies

*P. veronii* 1YdBTEX2 was grown on solid 21C mineral medium^39^ with toluene in the vapor phase at 30°C for 3 days. Colonies were inoculated into three 100 ml screw-cap conical flasks containing 25 ml of the mineral medium amended with 0.5 ml of a 1:19 mixture of toluene:tetradecane (Sigma-Aldrich ref: 34866; Aldrich ref: 87140). Flasks were incubated at 30°C and at 180 rpm on an orbital shaker until a culture turbidity (OD_600_) of 0.7 (mid-exponential growth phase). At this point, the three cultures were pooled, and the bacterial cells were harvested by centrifugation (swing-out rotor A-4-44, Eppendorf; 3220 g, 8 min, 30°C). The cell pellet was resuspended in the mineral medium and centrifuged again, as described above, to remove any residual carbon. After that, the cells were resuspended in mineral medium, diluted to obtain a starting OD_600_ of 0.16 and transferred into four replicate flasks, to which the toluene:tetradecane mix was added, as described above. The cultures were incubated at 30°C and 180 rpm and regularly sampled for OD measurements. For transcriptomics, approximately 1 x 10^9^ cells were harvested from an appropriate volume of the culture by centrifugation at 3,500 g, for 6 min at 30°C; snap-frozen in liquid nitrogen and stored at −80°C until RNA extraction. Three quadruplicate sample sets were produced: the inoculum at time 0 (T0h), that is cells starved for approximately 30 min at 25°C in mineral medium without a carbon source; exponentially growing cells (at OD_600_ ≈ 0.5, harvested after 4 h of incubation in the toluene-amended mineral medium); stationary phase cells (at OD_600_ ≈ 1.8–1.9 harvested after 24 h incubation). To study the composition of the exometabolome (metabolites in the culture medium excreted or leaked out of cells), 1 ml of the culture was sampled at the same time points. Cell culture was transferred into a 1.5 ml polypropylene microcentrifuge tube, which was clarified by 20 min centrifugation at 21,100 *g* and 4°C, after which 0.5 ml of spent media was transferred to a new polypropylene tube, stored at −80°C and shipped on dry ice to the metabolomics facility.

### RNA extraction, RNA-seq library preparation and sequencing

Total RNA from the frozen cell pellets was extracted using the RNA PowerSoil Total RNA Isolation Kit (MoBio Laboratories) as recommended by the manufacturer. Contaminating genomic DNA was removed by two cycles of TURBO DNase (Invitrogen) digestion and RNeasy MinElute Cleanup kit (QIAGEN) column purification. The quantity, purity and integrity of RNA samples were assessed using agarose gel electrophoresis, NanoDrop spectrophotometer (ThermoFisher Scientific) measurements and Agilent 2100 Bioanalyser (Agilent Technologies) profiling.

The completeness of DNA removal was verified by PCR using primers Pv_chr2_fw, ATCGGCTGTCCGACATCGGACG and Pv_chr2_rev, TCGAAGAGCTCCACCGAGAGCCGCC) and 1 pg of genomic DNA as a positive control, as described previously^23^. Next, 4 µg of each RNA sample were depleted from ribosomal RNAs, converted to the reverse-complement stranded Illumina sequencing library using the ScriptSeq Complete Kit (Bacteria, Illumina) and indexed with ScriptSeq™ Index PCR primers set 1 (Epicentre, Illumina) following the standard protocol. The resulting directional RNA-seq libraries were sequenced using single-end 100-nt read chemistry on an Illumina HiSeq 2500 platform (Illumina) at the Lausanne Genomic Technologies Facility.

### Untargeted LC-HRMS metabolomics

The spend media (100 µL) samples collected at different time points were extracted with 400 µL of ice-cold methanol to quench the metabolism, precipitate proteins and extract a broad range of polar metabolites. The extracted media were analyzed by HILIC-HRMS using an electrospray ionization source operating in both positive and negative mode. Pooled QC samples (representative of the entire sample set) were analyzed periodically (every 4 samples) throughout the entire analytical run in order to assess the quality of the data, correct the signal intensity drift and remove the peaks with poor reproducibility (CV > 30%) that can be considered chemical or bioinformatic noise. Data were acquired using a 1290 UHPLC system (Agilent Technologies) interfaced with a 6550 iFunnel Q-TOF mass spectrometer operating in a full-scan MS mode. In addition, pooled QC samples were analyzed in auto MS/MS mode (i.e. Data Dependent Analysis [DDA]) to acquire the MS/MS data for metabolite identification. In positive mode, chromatographic separation was carried out using an Acquity BEH Amide, 1.7 μm, 100 mm × 2.1 mm I.D. column (Waters, Massachusetts, US). The mobile phase was composed of A = 20 mM ammonium formate and 0.1% formic acid in water and B = 0.1% formic acid in acetonitrile. A linear gradient elution from 95% B (0–1.5 min) down to 45% B (17–19 min) was applied followed by 5 min for column re-equilibration to the initial gradient conditions. The flow rate was 400 μL/min, column temperature 30°C and sample injection volume 2 µl. ESI source conditions were set as follows: dry gas temperature 290°C and flow 14 L/min, fragmentor voltage 380 V, sheath gas temperature 350°C and flow 12 L/min, nozzle voltage 0 V, and capillary voltage 2000 V. In negative mode, a SeQuant ZIC-pHILIC (100 mm, 2.1 mm I.D. and 5 μm particle size; Merck, Damstadt, Germany) column was used. The mobile phase was composed of A = 20 mM ammonium acetate and 20 mM NH_4_OH in water at pH 9.3 and B = 100% acetonitrile. The linear gradient elution ran from 90% (0–1.5 min) to 50% B (8–11 min) down to 45% B (12–15 min). Finally, the initial chromatographic conditions were established during a 9 min post-run for column re-equilibration. The flow rate was 300 μL/min, column temperature 30°C and sample injection volume 2 µl. ESI source conditions were set as follows: dry gas temperature 290°C and flow 14 L/min, sheath gas temperature 350°C, nebulizer 45 psi and flow 12 L/min, nozzle voltage 0 V, and capillary voltage 2000 V.

In the MS-only mode, the instrument was set to acquire over the m/z range 50–1200, with the MS acquisition rate of two spectra/s. Targeted MS/MS data for dysregulated metabolite features were acquired using the inclusion list with narrow isolation window (≈ 1.3 *m/z*), MS acquisition rate of 500 ms, and MS/MS acquisition rate of 500 ms.

### Data processing and statistical analysis

#### Transcriptomics

Read mapping, sorting and formatting of the raw reads was done with Bowtie2^40^ and Samtools^41^, using the finalized gapless *P. veronii* 1YdBTEX2 genome sequence as in Morales et al^23^. Mapped reads were counted with HTSeq^42^, then further processed and analyzed with edgeR^43^. Only reads counted more than once per million in at least three replicates were kept. After normalization of the counts, transcript abundances were compared in pairwise conditions in a modified Fischer exact test (as implemented in edgeR). Genes were called significantly differentially expressed between two EXPO and STAT when their false-discovery rate was <0.05 and their fold-change >2 and were subsequently interpreted using Gene Ontology (GO) analysis. GO terms of *P. veronii* genes were inferred using the program BLAST2GO^44^. The same software was then used to analyze GO data sets of significantly differentially expressed genes in each pair-wise comparison, under the TopGO “Weight” algorithm.

#### Metabolomics

Raw LC-MS data were converted to mzXML files using ProteoWizard MS Convert. mzXML files were uploaded to XCMS for data processing including peak detection, retention time correction, profile alignment, and isotope annotation. Data were processed as a multi-group experiment, and the parameter settings were as follows: centWave algorithm for feature detection (Δm/z = 20 ppm, minimum peak width = 5 sec and maximum peak width = 30 sec, S/N threshold = 6, mzdiff = 0.01, integration method = 1, prefilter peaks = 3 prefilter intensity = 1000, noise filter = 0), obiwarp settings for retention time correction (profStep = 1), and parameters for chromatogram alignment, including mzwid = 0.015, minfrac = 0.5 and bw = 5.5. Preprocessed data (following the signal-intensity drift correction and noise removal with the “batchCorr” R package) were filtered according to the p-value (< 0.05) and signal intensity (> 1000 ion counts). The remaining table of metabolite features together with the most significant ion features selected from the loadings plot of the multivariate models in positive and negative ionization mode were subjected to metabolite identification as described below.

In the first instance, putative metabolite identification was performed by accurate mass and retention time (AMRT) matching against an in-house database (containing information on 600 polar metabolites from the Mass Spectrometry Metabolite Library Supplied by IROA Technologies, Sigma-Aldrich, characterized under the same analysis conditions). For that, raw data files (.d) were processed using Profinder B.08.00 software (Agilent Technologies) with the following parameter settings: mass tolerance 10 ppm, retention time tolerance 0.2 min, height filter 1000 counts, and peak spectrum obtained as an average of scans at 10% of the peak. In parallel, the XCMS output table of significantly different metabolite features was matched against the Human Metabolome Database (HMDB)^26^ based on accurate mass (AM) with Δppm = 10. The list of hits was further manually curated by taking into account the biological relevance of the hit (endogenous vs. exogenous metabolites) and a presence of the “true” peak shape (using the interactive XCMS Online interface^45^). Short listed ions of interest together with the metabolites identified by the in-house database were further subjected to targeted MS/MS validation. The metabolite identifications were validated by matching the MS/MS^46^ data acquired in the pooled samples (for each experiment in each LC-MS analysis mode) against the in-house PCDL database, METLIN (https://metlin.scripps.edu/) standard metabolite database^47^ or mzCloud (https://www.mzcloud.org/). Otherwise, if the MS/MS data quality did not allow for metabolite ID confirmation, the metabolite IDs remained putative, based only on accurate mass matching.

### Genome-scale model reconstruction

The *P. veronii* 1YdBTEX2 genome-scale metabolic reconstruction process combines the automated draft metabolic network process with several manual refinements and curation procedures, in total involving four main steps: **(i) functional annotation of the genome**: functional annotation of the genome is required prior to the reconstruction of the GEMs, and consequently, the quality of a GEM highly depends on the availability of a gapless genome. The protein sequences (FASTA files) of *P. veronii* 1YdBTEX2 were acquired from a previously published study^23^ and were annotated to identify the associated reactions that enzymes catalyze to determine the stoichiometric matrix, done using the RAVEN Toolbox^27^. The generation of the draft metabolic network followed the protocol detailed previously^27,48,49^, and the output of this annotation process is summarized in Table S6. Version 1.07 of the RAVEN Toolbox and the version of KEGG database as of September 2017 were used. **(ii) Compartmentalization, the definition of biomass reaction and uptakes and secretions:** In the absence of information or experimental evidence about the compartmentalisation and the cell content in *P. veronii* 1YdBTEX2, the cell compartment information (i.e., cytosol and periplasm), the transport mechanism between the compartments and the extracellular environment, the uptakes and secretions, and the biomass composition were obtained from two available models of closely related Pseudomonads*: Pseudomonas putida*^28–30^ and *Pseudomonas stutzeri*^31^. The biomass reaction in a GEM designates the metabolic precursors that build the BBBs, i.e., DNA, RNA, amino acids, lipids and carbohydrates and their corresponding stoichiometric coefficients. **(iii) Thermodynamic curation:** In a thermodynamically curated model, the standard Gibb’s free energy of a reaction and consequently the directionality of the reactions, i.e., reversible or irreversible, are associated with the reaction as additional constraints, which allow the performance of TFA. We followed the established protocol to thermodynamically curate metabolic models^32,33^ to add thermodynamics constraints to iPsvr, where we could estimate the standard Gibb’s free energy of formation of 76% metabolites, using the group contribution method (GCM)^50^ and the standard Gibb’s free energy of 84% of metabolic reactions in iPsvr. This allowed us to perform TFA, simulate the growth and determine whether iPsvr was functional, with further evaluations if it correctly predicted the expected growth-associate phenotypes. **(iv) Gap-filling of iPsvr:** The draft metabolic network of iPsvr did not contain all the necessary reactions for the production of all the biomass building blocks, and thus, the model did not show any growth. This is a very common observation in the reconstruction process of GEMs, since the (automatic) genome annotations are often incomplete or erroneous, with significant proportion of predicted proteins having no functions attributed^51^. Therefore, reactions without associated genes were included in order to obtain a functional model that simulates non-zero growth, done using a procedure called gap-filling. We followed the gap-filling procedure introduced previously^48^, wherein the production of each biomass precursor is defined as a metabolic task^27^ and a mixed-integer linear programming (MILP) formulation is used to generate alternative groups of minimal number of reactions (borrowed from KEGG) that enable the production of the biomass precursor. The draft GEM was gap-filled by iterative manual curation until a model was obtained that was able to carry non-zero flux through the biomass reactions at steady state (i.e., signifying ‘growth’). Although gap-filling is a routine procedure in the curation of GEMs and most of the available GEMs are gap-filled to obtain a functional model that is able to grow under defined conditions, gaps in pathways not involved with biomass production are, however, mostly overlooked in GEM analysis. We performed a complementary second gap-filling step apart from the biomass reaction, where as a result, the consistency of iPsvr with the obtained experimental data on gene expression and metabolomics was increased. Available GEMs usually contain a large well-connected subnetwork, which encompasses the most of the central carbon metabolism and a part of the secondary metabolism, and many isolated reactions (or sets of reactions) are probably disconnected from the rest of the network because of misannotations or insufficiently known pathways. Such isolated reactions/pathways are blocked, i.e., cannot carry flux under any condition, and therefore one or more reactions must be added to connect the blocked reactions with the rest of the metabolic network. To this end, the metabolic network structure of iPsvr was decomposed to the main component and to the isolated (disconnected) reactions/pathways using a MATLAB graph-based built-in function (conncomp). Using the same gap-filling approach, the blocked reactions/pathways that were associated with differentially expressed genes (in the pair-wise comparison of exponential- and stationary-phase datasets) or measured metabolites in exometabolomic data were gap-filled to become functional in the model (carry non-zero flux).

### Omics data integration

REMI^20^ was used for the integration of transcriptomics and exometabolomics data into iPsvr. REMI assumes that reaction fluxes associated with genes that are significantly differentially expressed are deregulated. Moreover, REMI also considers that the *in vivo* metabolite abundance ratios between the two conditions, e.g., the two growth phases can be used to constrain reaction fluxes associated with the metabolites that are differentially regulated. Expression/abundance-based ratios between the two conditions are formulated as flux perturbations for each reaction and are imposed as constraints on individual fluxes. Based on the data used, it translates into three different methods: (i) REMI-TGex allows the integration of relative gene expression data into a thermodynamically curated GEM, (ii) REMI-TM allows the integration of metabolomics data and (iii) REMI-TGexM integrates simultaneously both the gene expression and metabolite abundance data as additional constraints into the metabolic model.

REMI aims to maximize consistency between differential expression and fluxes as well as differential metabolite concentrations and fluxes. To study condition-specific differences in metabolism between two conditions (perturbed *vs* reference), REMI considers a separate metabolic model for each condition. Then, to integrate differential expression, REMI enforces a higher flux through a reaction in the perturbed condition (perturbed model) as compared to a reference condition (reference model) if the genes of the reaction are upregulated. For downregulated reactions, REMI enforces a lower flux as compared to a reference. To integrate extracellular metabolite concentrations, REMI assumes that if a metabolite is upregulated, then the production of the metabolite is forced to be higher as compared to a reference condition, and similarly, a lower production of a metabolite is forced if a metabolite is found to be downregulated. Then, an optimization problem is formulated to maximize the number of constraints imposed by the relative gene expression and metabolite abundances that can be integrated into the model while preserving a growth phenotype. Two scores are calculated: a theoretical maximum consistency score (TMCS), representing the number of genes/metabolites with available omics data, and the maximum consistency score (MCS), representing the number of genes/metabolites whose relative omics data are consistent with relative network fluxes and therefore can be integrated in the model.

The MILP formulation enables enumerating alternative sets (size equal to MCS) from a given set of constraints. The most consistent models are built by activating constraints that are overlapping and consistent between all the alternatives.

We used the transcriptomics and/or exometabolomics datasets measured at the exponential and stationary phase and also at the sand versus liquid environment, to derive additional flux constraints for the TFA problem, which were applied using REMI.

### Thermodynamics-based flux analysis (TFA) and the analysis of the biomass building blocks

To determine fluxes and subsequently the growth rate at different growth and environmental conditions using iPsvr, we employed FBA, the most widely used constraint-based modeling technique for studying biochemical networks and cellular physiology. Previous studies show that the integration of appropriate thermodynamic constraints leads to more accurate metabolic model predictions and also a significant reduction in the ranges of the predicted fluxes (solution space)^8,32^. We perform TFA for estimating the growth rate before and after the integration of omics data.

We further identified the biomass building blocks (BBB), wherein a low/zero production limits growth upon the transition of cells from the exponential to stationary phase or from liquid to the soil environment (two examples that were discussed in this work). Each BBB was tested by defining the TFA objective function as the maximum production of that metabolite under the defined media condition (the same for all the BBBs). Then, the BBB production was compared and the limiting BBBs for each case were identified.

## ACKNOWLEDGMENTS

This work was supported by SystemsX.ch, and evaluated by the Swiss National Science Foundation, within grant 2013/158 (Design and Systems Biology of Functional Microbial Landscapes ‘MicroScapesX’). The author would like to thank Dr. Ljubisa Miskovic for constructive criticism of the manuscript and for his careful proof-reading.

## SUPPLEMENTARY FIGURES

**Supplementary Figure 1:**
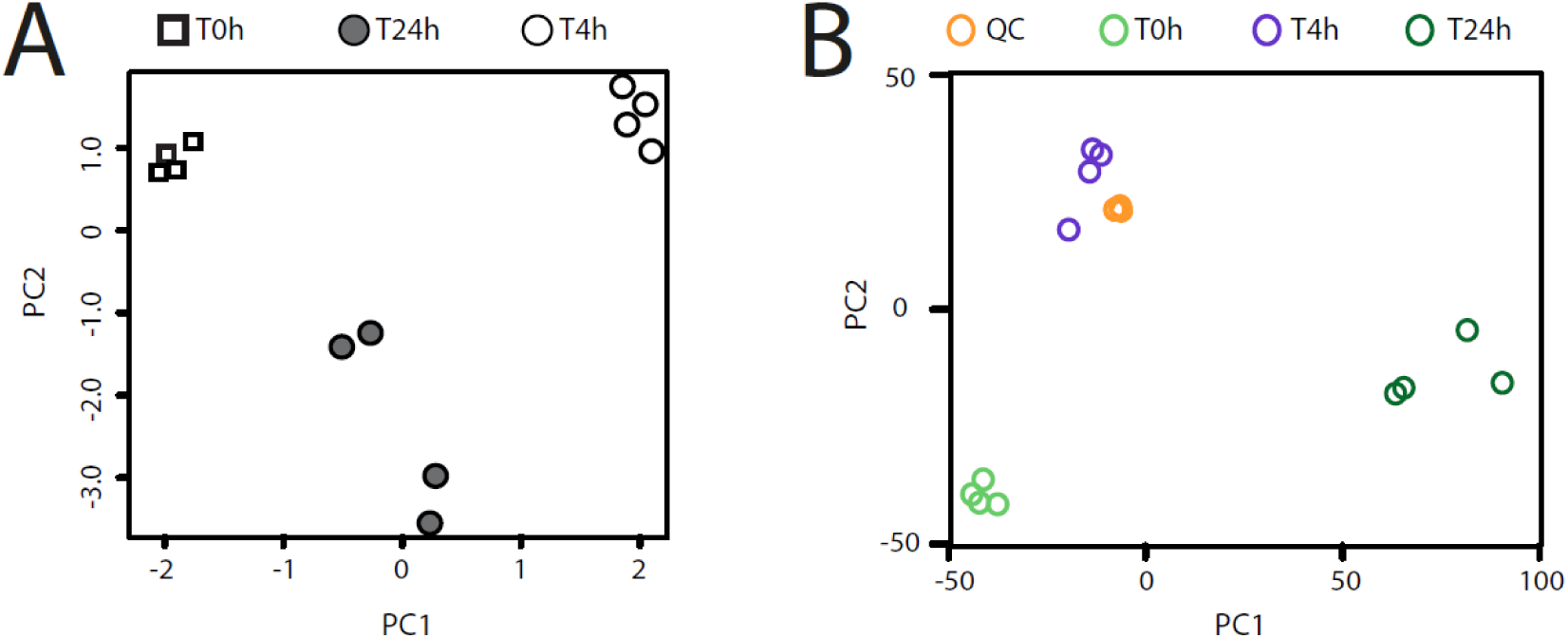
Principal component analysis of. (A) quadruplicate global RNA-sequencing data sets of *P. veronii* 1YdBTEX2 after 4 h, 24 h and the 0 h control (T0) and (B) of exometabolomics data, grouped QC samples of T0, T4 and T24 samples.

**Supplementary Figure 2:**
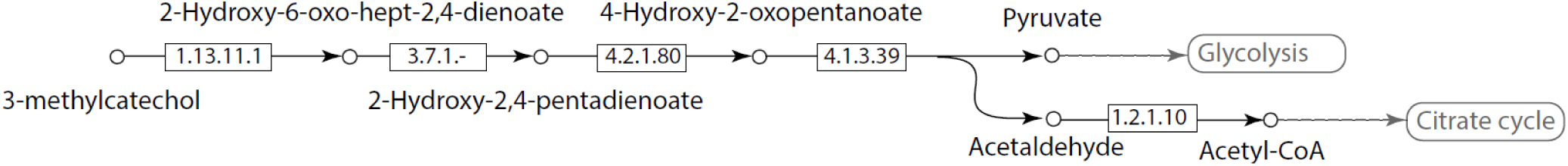
Main functional toluene degradation pathway branch in iPsvr. 3-methylcatechol is converted to pyruvate and acetaldehyde, which are further involved in central carbon pathways.

**Supplementary Figure 3:**
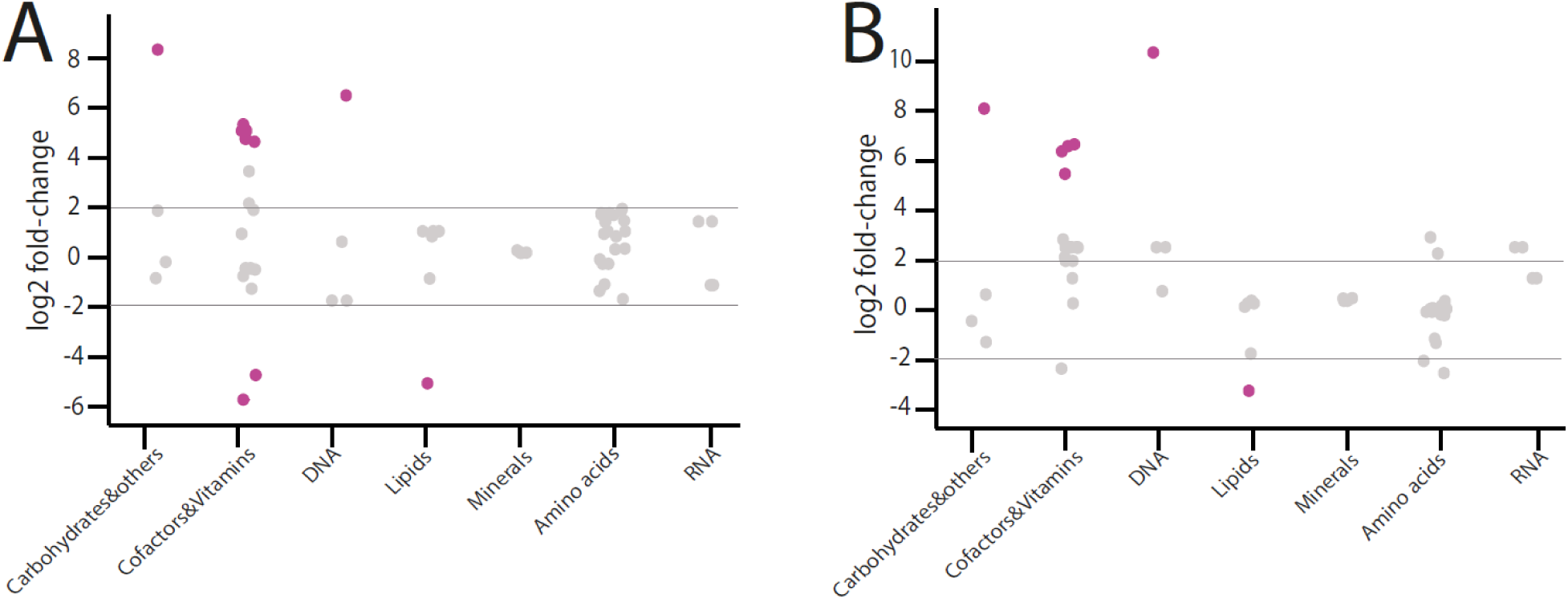
Differential *in silico* production (mmol/gDW/h) of seven biomass building block groups between cells in liquid medium and sand. (A) Integration of relative gene expression data (liquid vs sand) using REMI-TGex when cells grow on Toluene. (B) Integration of relative gene expression data (liquid vs sand) using REMI-TGex when cells grow on Succinate. Each dot in the graph represents the individual biomass precursors within that category and the statistically significant changes (p=value < 0.05) are highlighted in purple. For BBB group statistics, see Table S7.

## SUPPLEMENTARY TABLES (given as an excel file and each excel sheet is a Supplementary Table)

**Table S1.** Summary of RNA-seq yields of the different time points.

**Table S2.** Significantly differentially expressed metabolic genes in EXPO vs. STAT

**Table S3.** Under-represented Biological processes in STAT vs EXPO.

**Table S4.** iPsvr network decomposition into its main subnetwork and isolated reactions/pathways.

**Table S5.** Gap-filled reactions introduced into the iPsvr with their corresponding genes.

**Table S6.** Differential *in silico* production (mmol/gDW/h) of biomass precursors, grouped in seven biomass building block groups, between the exponential and stationary growth phase using REMI-TGexM, REMI-TGex and REMI-TM.

**Table S7.** Statistical test for BBB groups. Significant up- or down-regulations (p=value < 0.05) are highlighted in red.

